# Combining transcutaneous spinal stimulation and functional electrical stimulation increases force generated by lower limbs: When more is more

**DOI:** 10.1101/2023.12.22.573119

**Authors:** Alexander G. Steele, Albert H. Vette, Catherine Martin, Kei Masani, Dimitry G. Sayenko

## Abstract

**Background:** Transcutaneous Spinal Stimulation (TSS) has been shown to promote activation of the lower limb and trunk muscles and is being actively explored for improving the motor outcomes of people with neurological conditions. However, individual responses to TSS vary, and often the muscle responses are insufficient to produce enough force for self-supported standing. Functional electrical stimulation (FES) can activate individual muscles and assist in closing this functional gap, but it introduces questions regarding timing between modalities.

**Methods:** To assess the effects of TSS and FES on force generation, ten neurologically intact participants underwent (1) TSS only, (2) FES only, and (3) TSS + FES. TSS was delivered using four electrodes placed at T10–T11 through the L1–L2 intervertebral spaces simultaneously, while FES was delivered to the skin over the right knee extensors and plantarflexors. For all conditions, TSS and FES were delivered using three 0.5 ms biphasic square-wave pulses at 15 Hz. During the TSS + FES condition, timing between the two modalities was adjusted in increments of ¼ time between pulses (16.5 ms).

**Results:** When TSS preceded FES, a larger force production was observed. We also determined several changes in muscle activation amplitude at different relative stimulus intervals, which help characterize our finding and indicate the facilitating and inhibitory effects of the modalities.

**Conclusions:** Utilizing a delay ranging from 15 to 30 ms between stimuli resulted in higher mean force generation in both the knee and ankle joints, regardless of the selected FES location (Average; knee: 112.0%, ankle: 103.1%).

## Introduction

Spinal cord injury (SCI) is a life-long, devastating, and costly condition. Many individuals with SCI cannot stand by themselves due to paralysis of their leg muscles [1]. Affected individuals often spend a considerable amount of time in a seated position rather than engaging in standing activities, which has negative effects on mobility, autonomic functions, and quality of life [2]. Standing has various therapeutic benefits for individuals with SCI, such as improved blood circulation, respiration, bone density, skin health, and increased overall wellbeing [3]. Therefore, the standing posture has been used as a therapeutic approach for individuals with SCI (i.e., standing therapy) [4]. There are several therapeutic tools that enable individuals with SCI to stand upright, such as standing frames [5]. However, these tools provide only mechanical, i.e., “passive” support and do not require individuals to contract their leg muscles. The benefits of standing therapy are maximized during self-assisted standing [4], which can be accomplished with some external support such as walkers, knee-ankle-foot orthoses, or, more recently, powered exoskeletons, with the weight-bearing being achieved primarily by the legs [6]. This form of standing not only enhances muscle endurance and fatigue resistance but also facilitates active voluntary control of posture, such as during body-weight shifts. Considering the extensive reorganization of cortico-brainstem-spinal, corticospinal, and spinal networks following SCI [7], the functional improvement demonstrated during stand training will likely reflect the synergistic interactions within and between the brain and spinal cord, contributing to improved rehabilitation outcomes.

Neuromodulation approaches promoting physiological response in the lower limb and trunk muscles can be used in combination with activity-based therapies to potentiate functional outcomes. In this context, non-invasive Functional Electrical Stimulation (FES) which uses electrical pulses to excite the peripheral motor nerves to generate contractions of paralyzed muscles and can augment or induce specific movements [8-10]. However, the use of FES alone to promote standing has several disadvantages: first, FES needs to be delivered and controlled over multiple muscle groups [11]; second, peripheral neuromuscular stimulation, such as FES, inevitably results in fatigue as it recruits the fastest and most fatigable muscle fibers first [12]. Transcutaneous Spinal Stimulation (TSS), another non-invasive electrical stimulation technique, has recently received much attention from clinicians and researchers [13-16]. TSS can produce motor responses via Ia afferents – α-motoneuron synapses in multiple spinal segments, but also is capable to activate other neural components within the spinal cord, including interneurons, ascending sensory fibers in the dorsal columns, descending motor tracts, and other polysynaptic pathways [17-22]. Thus, TSS can activate multiple paralyzed muscles at once and induce self-assisted standing in individuals with SCI, while being delivered over the lumbar spinal cord [23]. Furthermore, findings from numerous studies indicate that TSS volleys, facilitated by synaptic projections, enable the physiological activation of lower limb muscles, effectively preventing muscle fatigue [12, 23, 24]. However, the individual responses to TSS vary greatly depending, for instance, on the strength of the lower limb muscles, neural excitability, or asymmetry due to SCI; therefore, TSS may not sufficiently drive bilateral muscle activation for self-assisted standing for some individuals [25]. In this light, we propose that: (1) by combining TSS and FES, we can achieve greater activation of leg muscles than using either of the neurostimulation approaches alone; and (2) by adjusting the timing of stimuli, we can maximize the stimulation effect. To quantitatively test these hypotheses, we separately examined the effects of combined TSS and FES on knee extensor and plantarflexion force generation in neurologically intact individuals using different relative timing schemes.

## Methods

### Participants

Ten adults with no known neuromuscular or musculoskeletal impairments (6 females 4 males; age 27.5 ± 6 years, height 167.6 ± 7.6 cm, body mass 64.8 ±13 kg) agreed to participate in this study and written informed consent was obtained from each of the participants. Experiments were conducted from July 2019 to December 2021 and the experimental procedures were approved by the Houston Methodist Research Institute institutional review board (Study ID: Pro00020081) in accordance with guidelines established in the Declaration of Helsinki. The study did not include minors or people from vulnerable populations.

### Experimental Setup

TSS was delivered to the skin over the lumbosacral spinal enlargement using an electronically triggered constant current stimulator DS8R (Digitimer Ltd., UK) via self-adhesive electrodes (PALS, Axelgaard Manufacturing Co. Ltd., USA). Four cathodes (diameter: 3.2 cm) were placed along the midline of the spine at the intervertebral spaces starting between the T10 and T11 and ending between the L1 and L2 vertebrae as seen in Fig. 1A. Two oval anodes (size: 7.5 cm x 13 cm) were symmetrically bilaterally placed on the abdomen. FES was delivered to the skin over the right knee extensors and right plantarflexors using a constant current stimulator DS8R (Digitimer Ltd., UK). The cathodes and anodes (size: 7.5 cm x 13 cm) were placed on the proximal and distal aspects of each muscle group, respectively. Both TSS and FES were delivered using a train of three 0.5 ms biphasic square-wave pulses at 15 Hz or with an inter-stimulus interval of 66 ms. Triplets were chosen to induce bursts, similar to functional stimulation delivery when fused muscle contractions promote movements [26].

**Figure 1.**
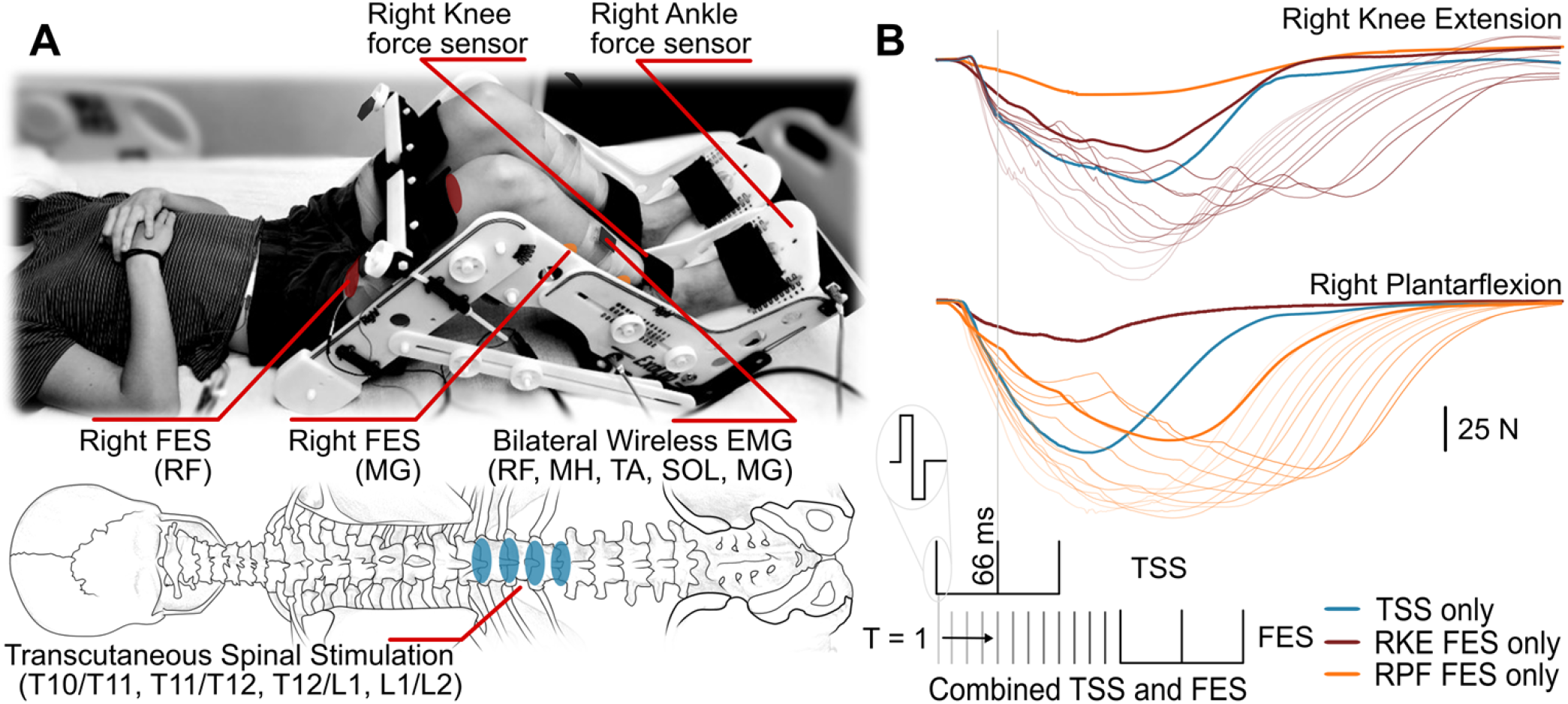
Experimental setup: (A) Transcutaneous spinal stimulation (TSS) was applied between the T10/T11 through L1/L2 spinous processes. Functional electrical stimulation (FES) was applied to the right knee extensors or plantarflexors. EMG was recorded bilaterally from the rectus femoris (RF) and medial gastrocnemius (MG) muscles. (B) Electronically controlled stimulation was delivered using biphasic triplet pulses with a 66 ms inter-pulse interval. TSS and FES were delivered at different relative stimulus intervals (RSI) ranging T = 1 to 180 ms, indicating TSS preceded FES and from -1 to -180 ms, indicating FES preceded TSS. Where RSI is the time delay of the first pulse in the FES triplet relative to the first pulse in the TSS triplet. Blue traces denote forces during TSS only, thick brown traces denote forces during right knee extensor (RKE) FES only, and thick orange traces denote forces during right plantarflexion (RPF) FES only. Note that while force sensors were independent for the knee and ankle joints, FES delivered to either knee extensors or plantarflexors inevitably caused changes in the measured force from synergistic joint. Shaded traces denote force response as RSI increases from RSI 1 (lightest) to RSI 180 (darkest) for TSS + FES of the right knee extensors (thin brown) and right plantarflexors (thin orange). Note that only forces generated by the FES targeted muscles are presented for FES + TSS.

Trigno Avanti wireless surface electromyography (EMG) electrodes (Delsys Inc., USA) were placed longitudinally over the left and right rectus femoris (RF) and medial gastrocnemius (MG) muscles. EMG data were amplified using a Trigno Avanti amplifier (Delsys Inc., USA).

The tests were performed in supine position with the participants’ legs placed in the Exolab apparatus (Antex Lab LLC, Russia) supporting the hip, knee, and ankle joints at 155, 90, and 90 degrees, respectively (Fig. 1A) [27, 28]. These joint angle positions were selected to isolate knee and ankle joint movements. Four calibrated, two-sided load cells (FSH00007, Futek, USA) measured forces generated by left and right knee extensors and plantarflexors independently. EMG and force signals were recorded at a sampling frequency of 2,000 Hz using a PowerLab data acquisition system (ADInstruments, Australia).

### Experimental Procedure

To determine motor threshold, which was defined as the first response > 2 S.D. of baseline recording, and maximum response, which was defined as two consecutive stimulation intensities where the mean peak-to-peak response did not change > 2 S.D. of baseline, TSS intensity was increased in increments of 5 mA starting at 30 mA. FES began at 5 mA and was increased in increments of 1 mA. Using an electronically triggered stimulator, three triplets were delivered at each intensity with a minimum of 2 seconds between triplets to reduce the effects of post activation depression, until the force generated by the knee extensors and plantarflexors plateaued or until maximum tolerated intensity. Then, TSS intensity for testing was selected to be approximately half the maximum force output of the TSS recruitment curves, and the FES intensity was selected to match the force output of the TSS intensity for the targeted joint.

Figure 1B outlines the experimental paradigm. The TSS + FES condition was performed in two blocks where TSS was paired with either FES of the knee extensors or plantarflexors. Before and after TSS + FES blocks, a control comprised of TSS only and FES only, with triplets delivered three times for each modality was performed. During TSS + FES, timing between the first TSS pulse and first FES pulse in the triplets was defined as the Relative Stimulation Interval (RSI) and varied using ¼ fractions of the 66 ms interstimulus intervals or 16.5 ms steps. Where RSI is defined as the delay the first pulse of the FES triplet was delivered relative to the first pulse of the TSS triplet. Each RSI was repeated three times. For simplicity of presenting the results are given as 15 to 180 ms RSIs in increments of 15. An RSI of -180 indicating that FES preceded TSS by 180 ms, and an RSI of 180 indicating that FES was delivered 180 ms after TSS.

### Analysis

Statistical analysis was performed using the open-source software RStudio (version 1.4.1717, RStudio PBC, Boston, MA). The integral of produced force by right knee extensors and plantarflexors was calculated at each RSI for both joints independently. Peak-to-peak EMG responses for the right RF and MG were calculated for the first response after TSS at RSIs -180, -165, -150, -60, -45, and -30. RSIs were selected to avoid the artifact caused by FES, which can obscure the response. For comparison across participants, data were normalized to TSS only for a given participant such that the average of pre- and post-TSS controls was 0. Data from each RSI were compared to the FES only applied at knee extensors or plantarflexors condition. For each comparison, normality was tested with a Shapiro–Wilk test. If the data were normally distributed (p > 0.05), a two-tailed paired Student’s t-test was used for analysis. If normality was violated, a Wilcoxon signed-rank test was performed. Significance was set at p < 0.05, and a Bonferroni correction was applied to account for multiple comparisons. Reported changes in forces are significant for the targeted joint (p < 0.05) unless otherwise denoted.

## Results

### Produced Forces

The time to peak produced force for TSS only occurred on average 205.5 ±19 ms and 200.0 ±15 ms (mean ± SD) after stimulation onset for the knee extension and plantarflexion, respectively. During FES of the knee extensors or plantarflexors, the average time to peak produced force was 208.6 ± 11 ms and 241.0 ± 5 ms after stimulation onset, respectively.

Figure 2 shows the mean and interquartile differences in the force-time integral during knee extension (Fig. 2A) and plantarflexion (Fig. 2B) forces during TSS + FES and FES only for each RSI, with 0% being the force produced from the TSS only control. For clarity triplet waveforms are shown above each plot for TSS (black) and FES (red) at differing RSI. During TSS + FES applied to the knee extensors (Fig. 2A), the first increase in produced knee force (blue) occurred during RSI 1 (knee: 98.8%, ankle: 32.1%). TSS preceding FES produced on average a higher mean force, the largest during RSI 105 (knee: 113.9%, ankle: 21.4%). When FES preceded TSS, the largest difference was during RSI -150 (knee: 76.1%, ankle: -2.5%).

**Figure 2.**
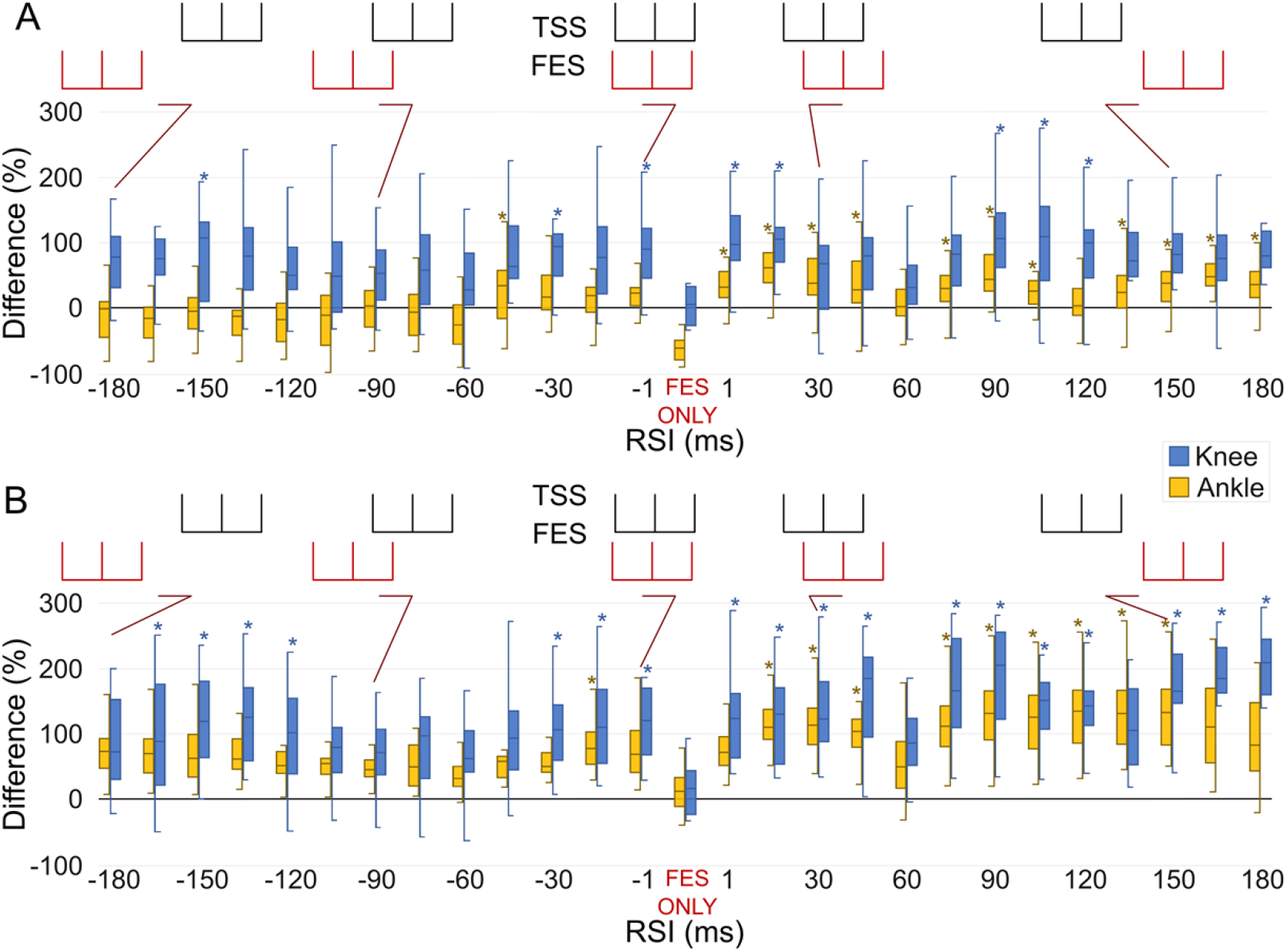
Produced forces: Difference in force-time integral measured as area under the curve, for the right knee extensors (blue) and right plantarflexors (yellow) during TSS + FES and FES only, as the relative stimulation interval (RSI) changes during (A) FES applied to the knee extensors and (B) FES applied to the plantarflexors. A negative RSI indicates that FES was delivered prior to TSS, and a positive RSI indicates that TSS was delivered prior to FES. Asterisks denote significant (p <0.05) changes compared to FES only. Data were normalized to average force produced by TSS only and are shown as the difference with respect to TSS only where TSS only is 0%. Triplet RSI are depicted above each plot for TSS (black) and FES (red) for several different RSI.

For FES applied to the plantarflexors (Fig. 2B), there was an increase in produced ankle force when TSS preceded FES at RSI 15 (knee: 133.0%, ankle: 110.3%) and when FES preceded TSS at RSI -15 (knee: 107.3%, ankle: 90.9%). As with knee extension, TSS preceding FES produced on average higher cumulative mean forces, with the largest difference at RSI 90 (knee: 204.1%, ankle: 138.7%). When FES preceded TSS, the only difference for the plantarflexors was during RSI -15 (knee: 107.3%, ankle: 90.9%).

### Spinally evoked motor potentials at RSIs of -60, -45, and -30

Figure 3 demonstrates evoked responses for RF and MG from a representative participant and grouped peak-to-peak data during TSS only and TSS preceded by FES. When FES was delivered at RSI -60 to knee extensors, the artifact obscured the RF response, but there was an 18.9% reduction of the TSS-induced response for MG when compared with RSI -45 (Fig. 3A). A similar effect was observed when FES was applied to the plantarflexors (Fig. 3B) where RF had a median reduction of 22.9% (p <0.01) when compared with RSI -45.

**Figure 3.**
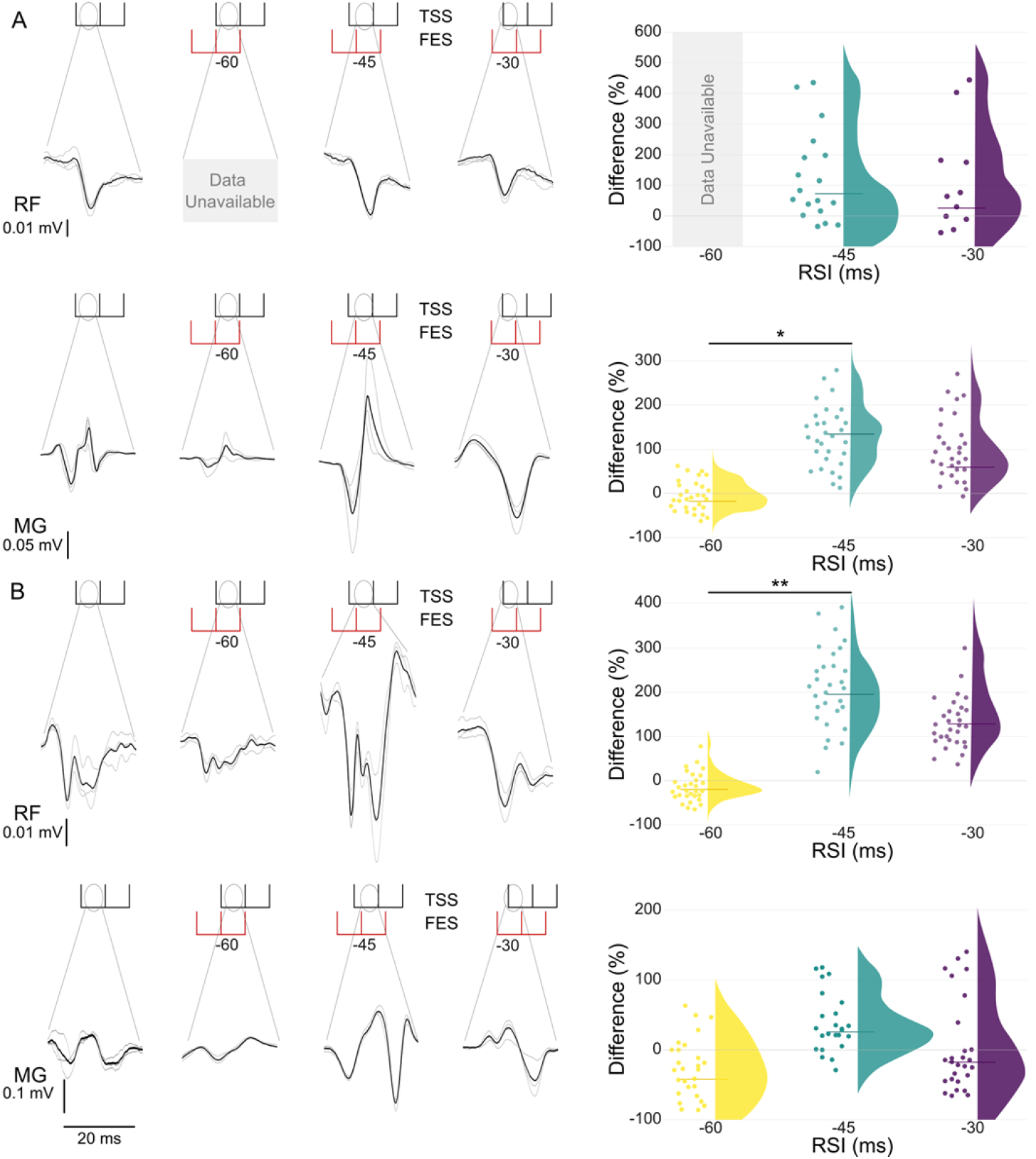
Changes in peak-to-peak amplitude of spinally evoked motor responses in right rectus femoris (RF) and right medial gastrocnemius (MG) during TSS only compared to TSS preceded by FES when (A) FES is applied to the knee extensors and when (B) FES is applied to the plantarflexors for a representative participant (left panels) and the response distribution of the group (right panels) for RSI -60, -45, and -30. Individual responses (left panels) are shown in light grey, with the average shown in black. Group data (right panels) utilize individual participant data points and a half violin with the line denoting the median. Data were normalized to average peak-to-peak response for TSS only and are given as the difference between TSS only and FES+TSS. Single and double asterisks denote significant (p < 0.05 and p <0.01, respectively) differences between RSI -45 and RIS -60.

### Spinally evoked motor potentials at RSIs of -180, -165, and -150

Figure 4 shows evoked responses for rectus femoris (RF) and medial gastrocnemius (MG) when TSS is preceded by FES for a representative participant (left) and group peak-to-peak responses (right). When FES was applied to knee extensors (Fig. 4A), responses were inhibited in RF and MG, with a 46.1%, 22.2%, and 37.7% median reduction for RF and a 58.4%, 61.8%, and 55.8% median reduction for MG during RSI’s -180, -165, and -150 respectively. When FES was applied to the plantarflexors (Fig. 4B), a 61.9% facilitation was found during RSI -165 when compared to TSS only.

**Figure 4.**
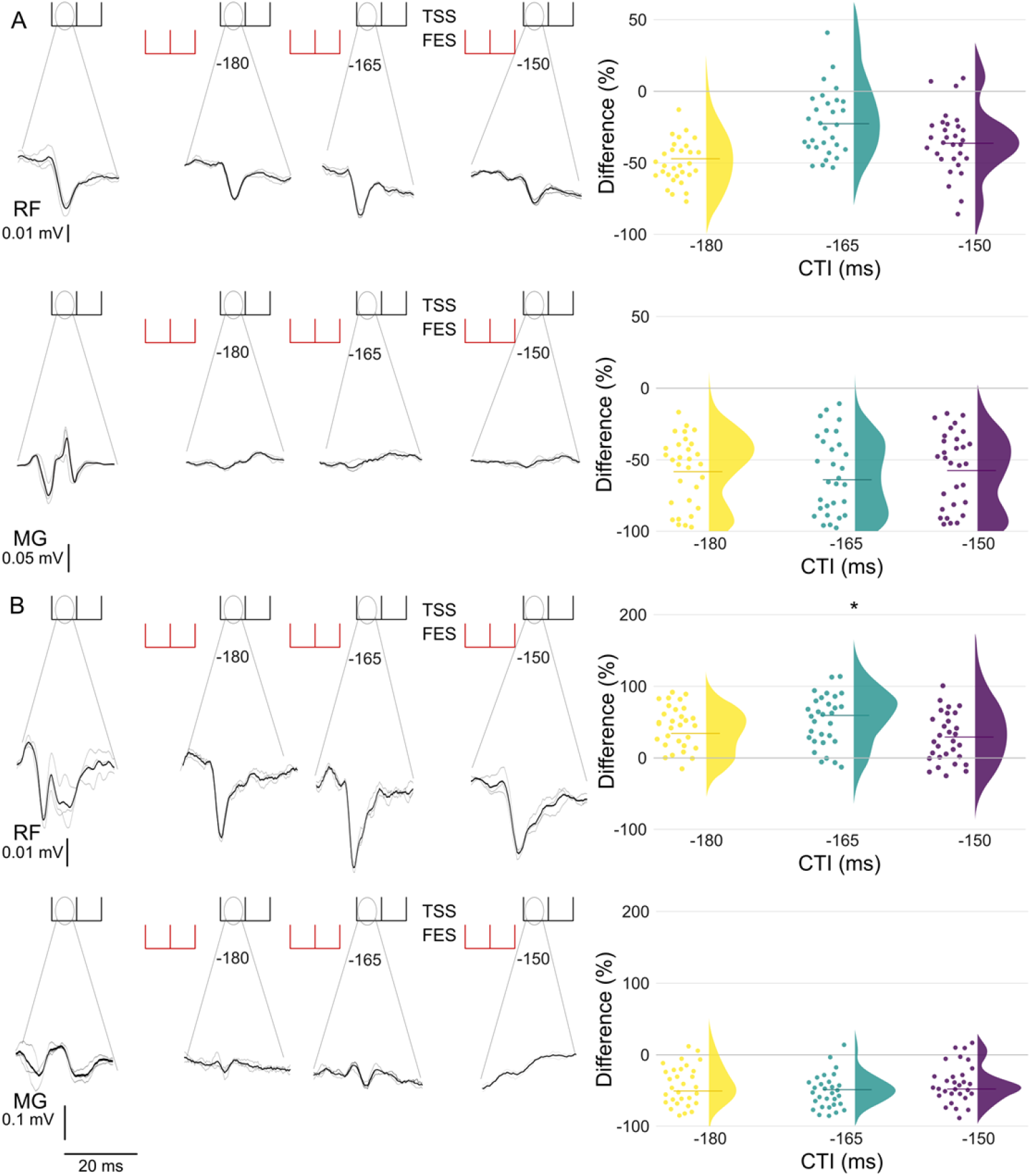
Changes in peak-to-peak amplitude of spinally evoked motor responses in right rectus femoris (RF) and right medial gastrocnemius (MG) when TSS preceded by FES for (A) FES applied to knee extensors and when (B) FES is applied to plantarflexors for a representative participant (left panels) and group data for RSI -180, -165, and -150 (right panels) shown utilizing individual points and a half violin where the line denotes the median. Data were normalized to average peak-to-peak response for TSS only and are given as the difference between TSS only and FES+TSS. Asterisks denote significant (p < 0.05) findings when compared to the TSS only condition.

## Discussion

To better understand the interaction between TSS and FES and their effect on force generation by leg muscles, we utilized TSS over the lumbar spine and FES over the knee extensors or plantarflexors separately and in combination, in neurologically intact individuals at rest. We demonstrated that TSS preceding FES resulted in larger produced force in both knee extensors and plantarflexors regardless of FES site across numerous RSIs, with the first occurrence of increased force as early as RSI 1. Interestingly, when FES preceded TSS (i.e., negative RSIs) there were only three instances of increased produced force in the knee joint during FES of knee extensors (RSIs -1, -30, and -150) and one instance of the increased force in the ankle joint during FES applied to plantarflexors (RSI -15).

When TSS preceded FES, the largest produced mean force occurred at RSI 90 regardless of FES site. This could be due to the time required for force generation. The average time to peak force production during TSS only was 205.5 ± 19 ms for knee extension and 200.0 ± 15 ms for plantarflexion, suggesting that timing the application of FES at the midpoint between the initiation of movement and peak produced force (i.e., approximately 100 ms) could increase force production due to the stimuli during mid-contraction. Alternatively, longer RSIs may reduce possible antidromic collision between FES and TSS. Other plausible mechanisms contributing to the observed increase in force-time integral when TSS precedes FES may involve TSS’s recruitment of smaller and less fatigable fibers initially or result from the interleaving of stimulus pulses, generating a net frequency twice that of each modality applied individually.

While the long-lasting artifacts from FES obscure the evoked response occurring at relatively short latency during positive RSIs for that muscle group, we observed the effect FES had on spinally evoked motor potentials by looking at changes during RSI’s -60, -45, and -30 (Fig. 3). We found inhibition (p < 0.01) at RSI -60 of the MG when FES was applied to the knee extensors and the RF (p < 0.05) when FES was applied to the plantarflexors when compared to RSI -45. This suggests that FES can interact with TSS in both an inhibitive and facilitating manner resulting in, based on the RSI, antidromic collision or potentiation on the presynaptic level.

When FES preceded TSS, there was almost no change in produced force for the targeted joint. However, FES applied to the plantarflexors prior to TSS (RSIs -1, -15, -30, -120, -135, -150, and -165) resulted in more instances of increased produced force in the knee joint than in the ankle joint. It is worth noting that analysis of the long RSIs (i.e., -180, -165, and -150) revealed inhibition for the RF and MG during FES applied to the knee extensors, but not when FES was applied to the plantarflexors. While similar levels of inhibition were found for the MG, there was no inhibition of the RF at the analyzed RSI’s and facilitation was found during RSI -165 (Fig. 4). This suggests that excitability of the motor pools projecting to the lower limb muscles can be modulated by the location of FES. This may be due to heteronymous Ia excitation produced by FES applied to the plantarflexors [29, 30]. Therefore, volleys from FES applied to the knee extensors converge at motor pools projecting to RF and MG inhibiting both RF and MG and FES applied to plantarflexors inhibits only MG and facilitates RF [30]. Further work should characterize this effect to determine the best ways for promoting this synergy, which may be beneficial for restoration of self-supported standing and walking in people with SCI.

From a functional perspective, the data suggest a 1 ms delay between the two modalities, while producing a higher force, is not the ideal case. Instead utilizing a delay of 15 to 30 ms between FES and TSS can yield more robust and synergistic response in knee extensors and plantarflexors, regardless of the selected FES site. Future work will characterize the interaction between FES and TSS in individuals with SCI.

## Limitations

Despite the insightful findings presented in this study, certain limitations warrant consideration. Firstly, the inclusion of neurologically intact participants, as opposed to individuals with SCI, may limit the generalizability of the results to the target population. Additionally, the experimental setup involved participants in a supine position rather than standing or engaging in other functional activities. While this choice was made to control variables, it does introduce a potential limitation regarding the translation of the observed effects to weight-bearing scenarios. Furthermore, the investigation into the mechanisms underlying the combination of FES and TSS was hindered by confounding artifacts in EMG data. These artifacts posed challenges in obtaining a detailed understanding of the intricate interactions between FES and TSS, highlighting the need for further research to delineate the specific mechanisms driving their combined effects.

## Conclusion

We demonstrated the interaction of TSS and FES on force generation and motor response in the leg muscles. We found that, when TSS preceded FES, a larger force production was observed, and that utilizing a delay between 15 to 30 ms between stimuli resulted in higher mean force generation by the knee extensors and plantarflexors regardless of FES location. Future work will be performed to determine if this effect is also seen in individuals with an SCI and to what extent it can be leveraged for self-supported standing during continuous stimulation.

## Data Availability

The datasets generated for this study are available from the corresponding author on reasonable request.

## Author Roles

### CRediT Author Statement

**Alexander G. Steele:** Investigation, Data curation, Visualization, Software, Formal analysis, Writing – Original Draft **Albert H. Vette:** Conceptualization, Methodology, Writing – Reviewing and Editing **Catherine Martin:** Investigation, Writing – Reviewing and Editing **Kei Masani:** Conceptualization, Methodology, Writing – Reviewing and Editing, Funding acquisition **Dimitry G. Sayenko:** Conceptualization, Methodology, Supervision, Resources, Formal analysis, Investigation, Writing – Reviewing and Editing, Funding acquisition

